# Genomic architecture of parallel ecological divergence: beyond a single environmental contrast

**DOI:** 10.1101/447854

**Authors:** Hernán E. Morales, Rui Faria, Kerstin Johannesson, Tomas Larsson, Marina Panova, Anja M. Westram, Roger K. Butlin

**Affiliations:** Centre for Marine Evolutionary Biology, Department of Marine Sciences, University of Gothenburg, Göteborg, Sweden; Department of Animal and Plant Sciences, University of Sheffield, Sheffield, UK; Centre for Marine Evolutionary Biology, Department of Marine Sciences at Tjärnö, University of Gothenburg, Strömstad, Sweden; IST Austria, Am Campus 1, 3400 Klosterneuburg, Austria

## Abstract

The genetic basis of parallel ecological divergence provides important clues to the operation of natural selection and the predictability of evolution. Many examples exist where binary environmental contrasts seem to drive parallel divergence. However, this simplified view can conceal important components of parallel divergence because environmental variation is often more complex. Here, we disentangle the genetic basis of parallel divergence across two axes of environmental differentiation (crab-predation vs. wave-action and low-shore vs. high-shore habitat contrasts) in the marine snail *Littorina saxatilis*, a well established natural system of parallel ecological divergence. We used whole-genome resequencing across multiple instances of these two environmental axes, at local and regional scales from Spain to Sweden. Overall, sharing of genetic differentiation is generally low but it is highly heterogeneous across the genome and increases at smaller spatial scales. We identified genomic regions, both overlapping and non-overlapping with recently described candidate chromosomal inversions, that are differentially involved in adaptation to each of the environmental axis. Thus, the evolution of parallel divergence in *L. saxatilis* is largely determined by the joint action of geography, history, genomic architecture and congruence between environmental axes. We argue that the maintenance of standing variation, perhaps as balanced polymorphism, and/or the re-distribution of adaptive variants via gene flow can facilitate parallel divergence in multiple directions as an adaptive response to heterogeneous environments.

## Introduction

Uncovering the evolutionary drivers of adaptive divergence is of central importance to understanding how biodiversity is generated and maintained (1, 2). Cases where phenotypic differentiation has emerged multiple times in response to similar environmental contrasts (i.e. parallel ecological divergence) represent ideal systems to study adaptive divergence (3). While cases of phenotypic parallel divergence are relatively common in nature (4, 5), it is not clear how often parallelism results from the same underlying genetic changes (6, 7). The expectation is for the genetic basis to be shared during parallel divergence, and for the amount of sharing to increase with decreasing evolutionary distance (according to meta-analysis; 7) and with decreasing geographic distance (according to modelling; 8). Importantly, different factors are likely to modify the amount of genetic sharing, including the congruence in direction for different environmental axes of parallel divergent selection across different locations (e.g. Fig. 1B), and the underlying genomic architecture (6, 9-11).

**Figure 1.**
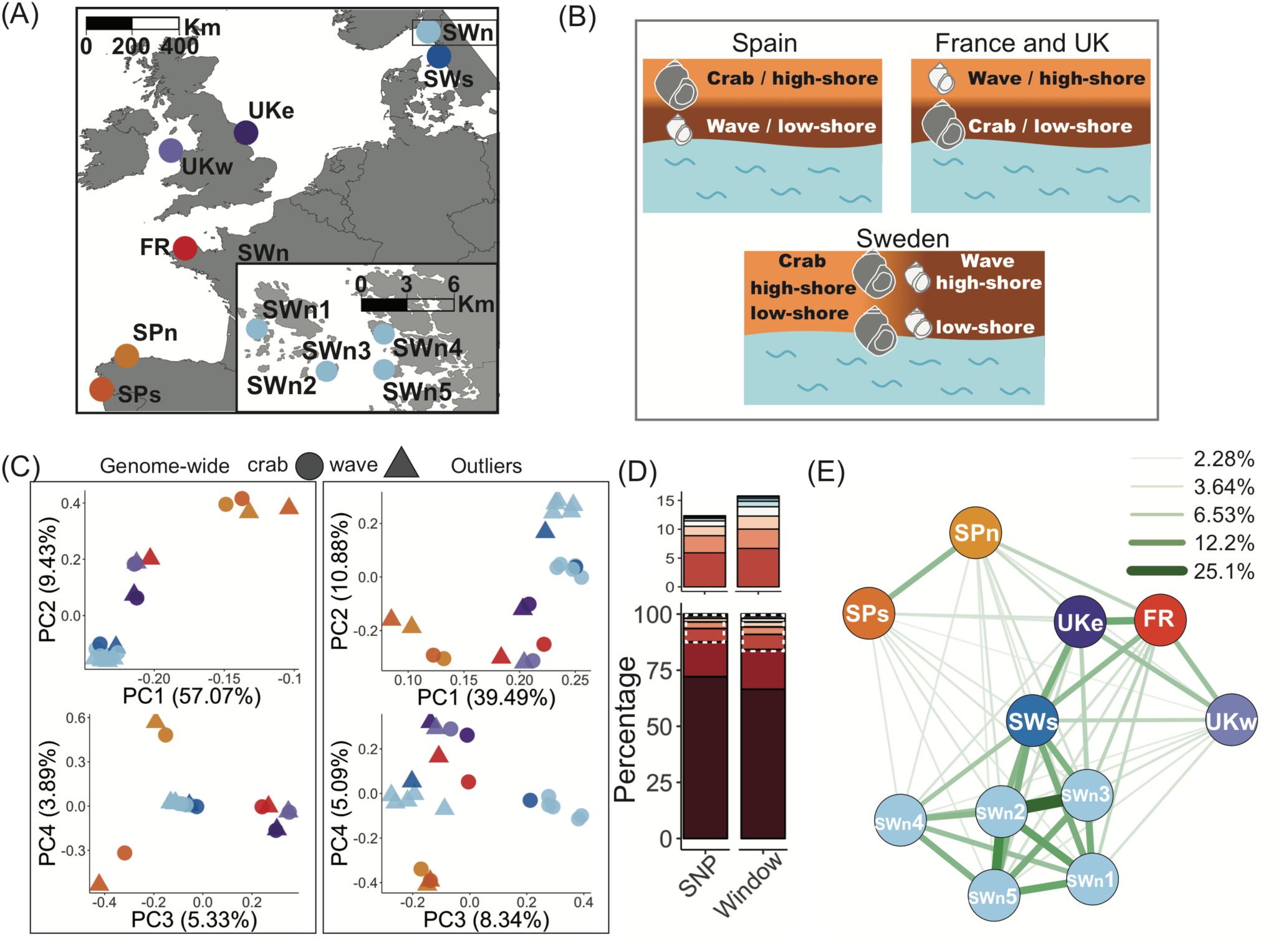
Samples, genetic structure and Crab-Wave genetic differentiation and sharing. (A) The 11 localities sampled at different geographical scales. SWn2 and SWn3 were sampled in different parts of the same island (< 1 Km). (B) Cartoon depicting the relative directions two axes of environmental divergence, Crab-Wave and Low-High, which are opposing between Spain and UK/France and orthogonal within Sweden (details in Supplementary Material). (C) PCA summarising genetic variation for 10,263,736 genome-wide SNPs (left panel) and for 705,786 highly Crab-Wave differentiated SNPs (i.e. outliers) identified across all localities (right panel). (D) Percentage of outlier sharing at SNP (N=705,786) and 500 bp window (N=22,315) levels. The colour ramp represents the number of localities that share outliers (from one [dark-red = unique outlier] to 11 [blue = fully shared outlier]), the dashed white rectangle shows a zoom-in of the top section. (E) Network plot showing the percentage of pairwise SNP outlier sharing between localities with thicker lines representing more sharing. The correlation between pairwise sharing and geographic distance was highly significant (Mantel r = −0.76, CI=-0.83/-0.66, P<0.001). The network plot of window-level sharing shows qualitatively similar results (see Fig. S7).

Even in presence of conspicuous phenotypic differentiation and replicated binomial habitat contrasts, it is important to consider additional axes of environmental and phenotypic differentiation (11, 12). Selective pressures that are unique, or act in different directions in each locality, are expected to lead to nonparallel patterns of phenotypic and genetic divergence (e.g. 10, 13). Thus, variability in environmental contrasts could influence sharing of genetic differentiation, but this has rarely been addressed directly (14, 15).

Genomic architecture is known to play a central role in parallel divergence (16, 17). In particular, chromosomal inversions, which suppress recombination in heterozygotes, may facilitate adaptive divergence by maintaining sets of co-adapted alleles (18-20). Inversions can act as reservoirs of adaptive standing variation and as vehicles to redistribute adaptive variation by gene flow, thus promoting parallel divergence with a shared genetic basis (21-24). On the other hand, inversions can confound inferences of the genetic basis of parallel divergence because they can modify apparent genetic sharing by capturing neutral loci within large haplotype blocks. Chromosomal rearrangements have been identified in studies of parallel evolution (e.g. 9, 25, 26), but the extent to which they contribute to shared genetic differentiation across multiple axes of parallel ecological divergence remains an open question.

Here, we assess how environmental variation and genomic architecture influence the genetic basis of parallel ecological divergence in the intertidal snail *Littorina saxatilis* (6, 27, 28). This species is broadly distributed across the north Atlantic and exhibits a series of life history traits conducive to the maintenance of adaptive divergence (reviewed in 27): Low dispersal due to restricted adult movement (29), internal fertilization and direct development (30), and the evolution of habitat choice (at least in Spain; 31). Large effective population sizes (high population densities, sperm storage and multiple paternity; 32) could lead to high rates of *de-novo* variation and to effective natural selection (29), including the maintenance of balanced polymorphisms observed in *L. saxatilis* (24, 33, 34).

*L. saxatilis* shows a strong pattern of parallel ecological divergence in at least two habitat contrasts. The main axis of divergence is between habitats dominated by crab-predation and wave-action. In areas of the shore where crab predation is high, *L. saxatilis* has evolved thick, large shells with a small aperture for the foot, and wary behaviour, as adaptations against crab predation. In areas of the shore exposed to strong wave energy, snails have evolved thin, small shells with a large aperture for the foot, and bold behaviour (6, 27). Here we refer to this axis simply as Crab-Wave divergence. The Crab and Wave habitats are adjoining and connected by narrow contact zones where gene flow occurs (34). Demographic modelling revealed that a scenario of multiple *in-situ* independent Crab-Wave divergence is a better fit to genome-wide neutral data than a scenario of ancestral divergence and secondary contact (28). A second axis of divergence is between low-shore and high-shore habitats. The Low and High environments experience contrasting thermal and desiccation conditions along a steep vertical gradient that impose strong selective pressures (35). Snails in high-shore habitats counteract higher desiccation exposure by lowering metabolic rates, exhibiting higher temperature resistance and lower water loss compared to snails in low-shore habitats (36-38). Here we refer to this axis simply as Low-High divergence. High-Low divergence has been observed in multiple locations but its demographic history has not been studied. The Crab-Wave environmental axis follows a different direction to the Low-High axis across multiple instances of parallel divergence (Fig. 1B; Supplementary Material). In Spain, the Crab habitat is associated with High-Shore and the Wave habitat with Low-Shore. In France and the UK, the direction is reversed. In Sweden, the two axes are orthogonal, each of the crab-predation and wave-action habitats contains a low-shore to high-shore gradient, with steeper environmental change in the wave-action than in the crab-predation habitat.

Seventeen candidate chromosomal inversions have been described in *L. saxatilis* from a single Swedish site (24; Fig. S1). Eleven of them showed a clinal pattern of allelic frequency change between Crab and Wave habitats (24). Three of these inversions correspond to genomic regions with clear signatures of Crab-Wave selection in a previous study from the same single Swedish site (34; see table S1 for details about all candidate inversions).However, the contribution of candidate inversions to parallel divergence is unknown.

We used the first whole-genome resequencing dataset for *L. saxatilis* to disentangle genomic regions associated with the two axes of ecological divergence across multiple localities from Spain to Sweden. We tested three general predictions:

I. Genome-wide sharing of genetic differentiation increases with geographical proximity. Populations with closer demographic and/or geographic links are expected to have a more similar genetic basis of parallel divergence (7, 8) due to stronger effects of gene flow and shared standing variation (22, 39).
II. Chromosomal inversion regions are enriched for shared outlier loci. If inversions contain adaptive standing variation and/or facilitate gene flow of adaptive variation they should promote the similar patterns of genetic divergence across multiple instances of parallel divergence (20, 23).
III. Genetic differentiation may be influenced by more than a single environmental axis of divergence. Here, we quantify the direction of genetic differentiation across each of the two axes, Crab-Wave and Low-High, and test how well chromosomal inversions explain differentiation in each case.

Testing these predictions simultaneously provides new insight into the impacts of geography, environmental variability and genomic architecture on the repeatability of evolution.

## Results

We performed genome resequencing of 1,744 individuals pooled into 26 pool-seq libraries from 11 northern European localities using a hierarchical design covering local and regional scales (Fig. 1A; Table S2). In Sweden, the shore level contrast was sampled within Wave and Crab habitats only in SWn3 and SWn5. Thus, the number of pools used in Crab-Wave analyses was 22 while 18 pools were used in Low-High comparisons (see details in Supplementary Methods; Table S2). Pool-seq is cost-efficient and recovers accurate population-level allelic frequencies (40), but can be biased when calculating differentiation metrics as it is not possible to distinguish the source of each sequencing read (41). Our approach was robust because: (i) we used a large number of individuals per pool (most pools contained 100 individuals; Table S2); (ii) we sequenced pools at high depth (mean = 68X); (iii) pool-seq allelic frequencies were highly correlated with those from individual-based sequencing from one locality (r^2^ > 0.88; Fig. S2); (iv) our genetic differentiation estimators were highly correlated (r^2^ = 0.95; Fig. S3) with recently developed alternative metrics (41); and (v) the outlier loci identified by these methods overlapped strongly (mean = 92%; Fig. S4).

### Sharing of Crab-Wave genetic differentiation is low but increases with geographical proximity

Localities varied in their Crab-Wave genome-wide genetic differentiation (ranges of mean F_ST_ in 500 bp non-overlapping windows: Spain = 0.09-0.12; France = 0.03; UK = 0.01-0.03; Sweden = 0.05-0.07; Fig. S5). We summarised genome-wide genetic variation with a PCA of 10,263,736 bi-allelic SNPs (Minor Allele Frequency, MAF > 5% in at least one pool). The first four PCA axes, containing most of the variation (66.5%), depicted a genetic structure consistent with geography (Fig. 1C). This genetic structure agrees with the previously-inferred *L. saxatilis* biogeographic history: populations in UK, France and Sweden originate from a different glacial refuge from Spanish snails, and our Spanish sites are separated by a north-south phylogeographic discontinuity (28, 42-44).

We identified Crab-Wave F_ST_ outliers as those within the top 1% quantile in each of the localities at two different levels: 705,786 outliers at the SNP-level and 22,315 outliers at the 500 bp window-level. In a PCA with all outlier SNPs, the first four axes (63.8% of variation) depicted a hierarchical structure of strong differentiation between countries and weaker ecotype differentiation within countries, suggesting low levels of overall outlier sharing (Fig. 1C; Fig. S6). Next, we directly measured the amount of outlier sharing across all localities. Most of the SNP- and window-level outlier loci (>66%) were unique to their locality. However, more than 15% of outliers were shared by at least two localities (Fig. 1D). Although only a small percentage of outliers was shared across all localities (<0.016%;), we would not expect a single fully shared outlier by chance. Higher sharing was observed at the window-level than at the SNP-level (Fig. 1D), indicating that shared genomic regions often contain different divergent SNPs across localities. Genome-wide, pairwise SNP outlier sharing varied between 2.3% and 25% across geographic regions (Fig. 1E), always higher than the random expectation of sharing between two localities (1%, 99% CI [0.9, 1.07]; see Supplementary Methods). The pattern of outlier sharing had a strong geographical signal where nearby localities shared a higher number of outliers than localities further apart. Previous genome scans focusing on small portions of the genome have shown a similar pattern where Crab-Wave outliers are rarely shared between regions or countries (44-48), but outlier sharing increases at nearby localities (39). Thus, we found support for our prediction 1 that genome-wide sharing of genetic differentiation is generally low but increases with geographical proximity.

### Genomic clusters of Crab-Wave differentiation coincide with putative chromosomal inversions

Using a recently developed linkage map for *L. saxatilis* (34), we were able to place half of the total genome content into 17 linkage groups (LGs). The map resolution is moderate (∼0.5 cM), so multiple scaffolds are associated with the same map position. The patterns of outlier sharing did not vary between scaffolds placed in the linkage map and those not placed (Fig. S8). We observed a heterogeneous landscape of genomic differentiation between ecotypes (mean Fst in Fig. 2A; FST per locality in Fig. S9), as commonly observed in other natural systems when divergence proceeds in the face of gene flow (e.g. 49, 50). On average, pairwise outlier sharing across the genome was relatively low (mean = 10.6% SD = 8.5; Fig. S10) in agreement with previous studies that focused on small portions of the genome (44-48). However, it was also highly variable, reaching far larger values in some LGs (e.g. in LG6, mean = 24.5% SD = 16%; Fig. S10).

**Figure 2.**
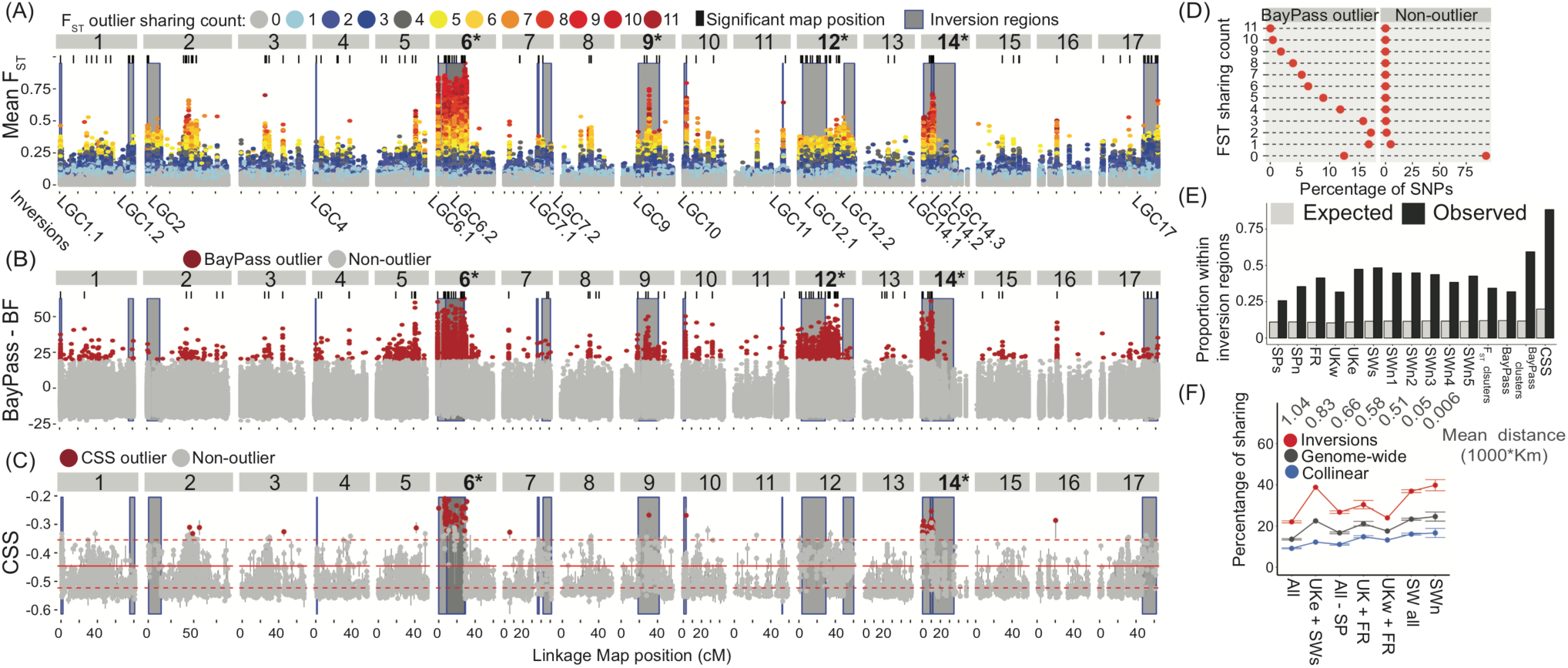
Genomic landscape of parallel Crab-Wave divergence, sharing and the influence of chromosomal inversions. (A) Average Fst between ecotypes across all localities, by linkage map position. Dots are coloured according to their outlier sharing count. (B) BayPass covariate model. Outlier SNPs (red) were defined by BF-score > 20. (C) Cluster Separation Score (CSS) per map position with their 95% confidence intervals (100 bootstrap). The red lines represent the mean and 95% CI for a genome-wide permutation. Outlier map positions (red) are defined by non-overlapping CIs. In panels A-C, LG names in bold and with an asterisk indicate a significantly higher outlier count compared to genome-wide random permutation, and black bars represent map positions with significantly higher counts compared to a random permutation within the same LG (Pval < 0.01). (D) The percentage of BayPass outliers and non-outlier SNPs that fall in each of the Fst outlier sharing count categories. (E) Expected vs. observed count of outliers within inversion regions for Fst outliers for each locality, significant Fst map positions (i.e. black bars in panel A), BayPass outliers, significant BayPass map positions (i.e. black bars in panel B), and CSS outliers. (F) Average pairwise sharing of F_ST_ outliers and average pairwise geographic distance across different comparisons.

We used three different methods to investigate Crab-Wave differentiation across the genome (see Supplementary Methods for details). (i) The sharing of outlier SNPs (top 1% F_ST_) across localities (as defined above). (ii) The covariate model from BayPass (51) to measure the association between allele counts and Crab vs. Wave habitat membership per SNP across all localities, while accounting for shared demographic history with the covariance matrix, Ω (52; Fig. S11). (iii) The cluster separation score (CSS) as in (9) to measure the strength of Crab-Wave differentiation, relative to that between localities within Crab and Wave habitats, at the level of map positions. The higher the CSS, the greater and the more consistent the Crab-Wave differentiation was across localities. Next, we tested for clustering of highly differentiated loci at two genomic scales (using values drawn from each of these three measures; see Supplementary Methods for details). At the LG scale, we identified significant LG values as those higher than the 95th percentile of a random genome-wide distribution, while at the map position scale, we identified significant map position values as those higher than the 95th percentile of a random distribution within the same LG.

Overall, we found that LG6, LG9, LG12 and LG14 showed significantly higher and more consistent than expected levels of Crab-Wave differentiation (F_ST_ and CSS) and covariation with the Crab-Wave axis (BayPass) across localities (significant LGs for each method are highlighted in bold in Fig. 2A-C). Nevertheless, some map positions with significantly higher than expected levels of shared Crab-Wave differentiation/covariation were scattered throughout the genome (significant map positions are highlighted with black bars in Fig. 2A-B; red dots in Fig. 2C). Some of the LGs harbouring significant clusters showed relatively low mean F_ST_ values (e.g. LG17) when compared to other parts of the genome (e.g. LG6) (Fig. 2A), suggesting that outlier sharing in some genomic regions is restricted to fewer localities or involves weaker differentiation.

To further investigate the geographic pattern of shared Crab-Wave differentiation across the genome, we tested pairwise F_ST_ outlier sharing within smaller geographic areas (e.g. within Spain or UK + France). For example, sharing in the LG17 cluster (around position 60 cM) was strong in Spain and less prevalent elsewhere, while strong sharing in LG2 (around position 50 cM) was limited to comparisons involving Spanish and Swedish localities (see Fig. S12 for a genome-wide overview). This suggested that some SNPs with Crab-Wave habitat associations identified with BayPass may not always be accompanied by elevated levels of Crab-Wave F_ST_. To confirm this, we compared BayPass BF outliers to the F_ST_ outlier sharing count. Most of the F_ST_ outlier SNPs (98.7%) identified (in any locality) were not identified as BayPass outliers. Moreover, among BayPass outliers, 12% were not identified as F_ST_ outliers at all, and of those that were, 49% had low F_ST_ outlier sharing counts (shared by 3 localities or fewer; Fig. 2D).

Lastly, we used the coordinates of the candidate inversions (24) identified at a single Swedish location (SWn4), to investigate the contribution of genetic differentiation within these regions to the geographical pattern of shared ecotype differentiation across localities. We refer to these genomic regions as ‘inversion regions’ (and other parts of the genome as ‘collinear regions’) but stress that polymorphic inversions have only been demonstrated at one site (SWn3; Fig. S1; Table S1). We found that inversion regions always contained a higher proportion of outliers, compared to the random expectation defined by the proportion of the genome that they contain (Fig. 2E; chi square P-val < 0.001). Next, we measured the level of pairwise F_ST_ outlier sharing within inversion regions and collinear regions at different spatial scales to ask whether sharing increases at smaller scales (Fig. 2F). Regardless of which genomic regions were considered, sharing increased with geographical proximity, except for a few comparisons that deviate from this trend. Notably, inversion regions always contained a higher percentage of sharing than collinear regions, suggesting that some inversions are important for Crab-Wave divergence across all localities. The findings in this section support our prediction 2: many outlier loci for Crab-Wave differentiation were strongly clustered across the genome and the clustering pattern was partly explained by the presence of putative chromosomal inversion regions.

### Genomic differentiation varies according to different axes of environmental variation

Populations undergoing parallel ecological divergence can experience unique local environmental conditions that could drive divergence in multiple directions (11). Apart from the Crab-Wave axis, *L. saxatilis* experiences a second axis of Low-High ecological variation (35, 38). Here we contrast these two axes of selection, exploiting their varying directions across different localities (see Introduction; Fig. 1B; and Supplementary Material).

To identify genomic regions that are related to the effect of Low-High divergence, we repeated the BayPass and CSS analyses and the clustering analyses at the LG and map position scales, this time focusing on the Low-High axis. Both BayPass and CSS analyses identified several LGs with clusters of differentiation consistent with strong parallel Low-High divergence (Fig. 3A-B). LG9, LG12 and LG14 had a significant over-representation of BayPass outliers compared to a genome-wide random permutation (Fig. 3A) and CSS showed similar results except that LG10 was highlighted and LG14 was not (Fig. 3B). Some of the significant clusters from the BayPass analysis appeared in genomic regions similar to those observed for the Crab-Wave divergence. A direct comparison of BayPass BF scores revealed a weak correlation between the two axes (r^2^=0.16, P<0.001), as 87% of outlier SNPs were found along the Crab-Wave divergence axis, 11.7% along the Low-High divergence axis and only 1.3% were implicated on both axes (Fig. 3D).

**Figure 3.**
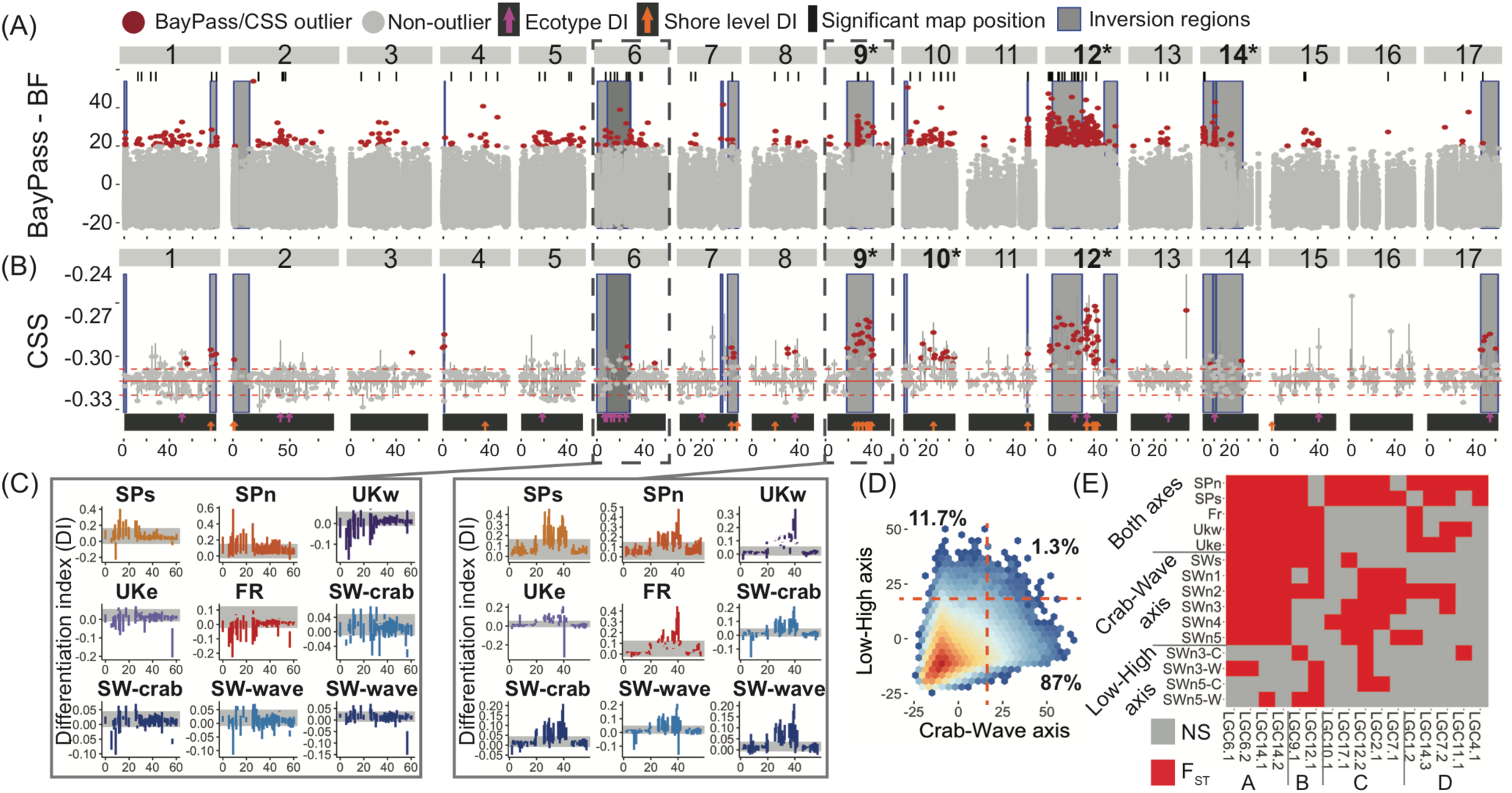
Two axes of parallel ecological divergence, Crab-Wave vs. Low-High habitat contrasts. (A) BayPass covariate model for Low-High divergence. (B) Cluster Separation Score (CSS) for Low-High divergence. Purple vs. orange arrows show map positions that were classified as differentiated on Crab-Wave or Low-High axes of divergence, respectively, according to their differentiation index (DI; see main text for expectations). (C) The coloured lines show the DI values for each locality. Rectangles of grey shading represent ±1.5 standard deviations of the genome-wide DI mean value for a given population, the threshold used to classify positions as ecotype or shore level divergence (purple vs. orange arrows in panel B, see text). Here we show plots for two LGs as examples, see Fig. S13 for others. (D) 2D-density plot comparing BayPass BF scores between Crab-Wave and Low-High tests. The grid colours show the range of count values from blue to red to reflect density of overlapping points. Orange dashed lines represent the BF-score = 20 thresholds that define each group with their percentages indicated. (E) Heatmap showing in red inversion regions that have significantly higher mean Fst compared to the 99% CI of the collinear genomic background. Non-significant (NS) tests are indicated in grey. Genetic differentiation (Fst) was measured per locality according to their direction along Crab-Wave and Low-High axes as indicated on the y-axis (see main text; Fig. 1B).

A problem with F_ST_ as a measure of differentiation is that it lacks directionality, potentially interfering with interpretation of outlier sharing (39). Therefore, we developed a metric, the differentiation index (DI), to measure and compare the strength and direction of allelic frequency differentiation on each axis of divergence. The DI is a normalized measure of Low-High genetic differentiation at a locality, using the two Spanish pools to confer directionality in assigning reference alleles (see Supplementary Methods). Given that the two axes of environmental divergence follow different directions among localities, we expect the DI to depend on whether a given genomic map position is implicated in Crab-Wave or Low-High divergence as follows: (1) Crab-Wave divergence: positive in Spain, negative in France/UK, and close to zero in Sweden. (2) Low-High divergence: positive across all localities. The DI analysis confirmed that the two axes of ecological divergence have different effects across the genome. For example, LG6 showed DI directions consistent with Crab-Wave divergence, while LG9 instead fitted the expected DI directions for Low-High divergence (Fig. 3C; see Fig. S13 for the remaining DI plots).

We applied the DI test to all map positions across the genome, asking whether DI values met the expected directionality (see above) in at least one locality within each of the three groups (Spain, France/UK and Sweden). The DI values of most map positions (96.3%) did not meet the expected directions for either Crab-Wave or Low-High divergence. We found 21 map positions that were consistent with the expectations for Crab-Wave divergence, and 26 that were consistent with the expectations for Low-High divergence (purple and orange arrows in Fig. 3B). While some of these map positions are scattered across the genome, some form clusters that often coincide with inversion regions (Fig. 3B).

### Chromosomal inversion regions are differentially involved in two axes of parallel divergence

Next, we asked whether mean F_ST_ within each of the inversion regions was significantly higher than the 99% confidence interval of F_ST_ at genome-wide putative collinear regions in each locality (see details in Supplementary Methods). Our findings indicated that some chromosomal inversions are likely involved in the Crab-Wave divergence and others in the High-Low divergence. Specifically, in line with our other results, we observed that inversion regions in LG6 and LG14 (group A in Fig. 3E) were universally involved in Crab-Wave divergence as they were significantly more divergent in all of the expected localities. Likewise, inversion regions in LG9 and LG12 were involved in Low-High divergence, as they were significantly more divergent in most of the expected localities (group B in Fig. 3E). The patterns observed for other inversions are more complex, some are predominantly differentiated in Spain and Sweden (group C in Fig. 3E), while others show no clear pattern (group D in Fig. 3E).

Overall, the results support our prediction 3: accounting for the direction of both Crab-Wave and Low-High axes of divergence is essential to understand the genomic basis of parallel ecological divergence in *L. saxatilis*.

## Discussion

Investigating the genetic basis of parallel ecological divergence is a key step towards understanding if, and to what extent, evolution is predictable (3-6). Here, we studied patterns of shared genomic divergence in one of the best-established natural systems of parallel ecological divergence, the marine snail *Littorina saxatilis* (6, 28). While previous studies focused on a single axis of ecological divergence in *L. saxatilis*, here we disentangle patterns of genome-wide differentiation across two major axes of parallel ecological selection: Crab-Wave and High-Low divergence. Specifically, we performed the first whole-genome study of genetic differentiation along each of these ecological axes of divergence and measured the explanatory power of the underlying genetic architecture. Overall, genomic differentiation can be explained by geographical proximity, the presence of putative chromosomal inversions and by accounting for the effects of multiple environmental axes, supporting our predictions.

### Adaptive substrate for parallel divergence via demographic and geographic links

Our results indicate that accounting for the demographic and geographical context of genetic differentiation and its underlying genetic architecture is essential to understand the evolution of parallel ecological divergence. Specifically, we showed that while sharing of outlier loci is generally low, sharing increases with geographical proximity. This finding indicates that localities with closer demographic histories likely share more standing variation and, at the same time, gene flow can more easily spread adaptive variation between geographically closer localities (1, 3, 22, 39). Levels of shared genetic differentiation were highly heterogeneous across the genome, with a few genomic regions and LGs accounting for most of the sharing. Notably, we were able to differentiate genomic regions, both inside and outside putative inverted regions, that are involved in parallel Crab-Wave divergence from those involved in parallel Low-High divergence. This finding suggests that *L. saxatilis* was able to repeatedly use reservoirs of standing genetic variation and/or source genetic variation via gene flow in order to evolve locally adaptive traits in heterogeneous environments.

### Chromosomal inversions as reservoirs and vehicles for parallel divergence

Outlier sharing is clearly magnified within putative chromosomal inversions that here we defined on the basis of polymorphisms at a single Swedish site (24). Such regions might contain loci that have been repeatedly used as the adaptive substrate for parallel ecological divergence and are involved in the establishment of strong barriers to gene flow by suppressing recombination across large genomic regions (21). Presumably they also contain many neutral loci with high divergence as a result of association with adaptive loci. Parallel divergence in *L. saxatilis* is believed to have occurred recently at higher latitudes, after postglacial colonization (28), but the exact role of putative chromosomal inversions during rapid parallel divergence in *L. saxatilis* is not clear yet. We found strong evidence of widespread chromosomal inversions differentially involved in two axes of ecological divergence, parallel Crab-Wave divergence (on LG6 and LG14) and parallel Low-High divergence (on LG12 and LG9). One plausible scenario is that most of the inversion polymorphisms evolved long before the last glaciations, possibly contributing to divergence in the distant past, and more recently fuelled rapid parallel divergence (24, 53-55). Further work is needed to confirm the presence of chromosomal inversions across the species range and to clarify the geographically complex patterns of differentiation of some inversions. Some of the putative chromosomal inversions may have recent origins or, alternatively, inversions could be variable in their allelic content and associated fitness effects.

Chromosomal inversions are known to store ancestral polymorphism that can be used later as a substrate for adaptive divergence and can be easily distributed via gene flow as “adaptive cassettes” (23, 56). Chromosomal inversions acting as polymorphism reservoirs is in line with our findings and those of Westram et al. (34) and Faria et al. (24), who used a densely sampled transect across a hybrid zone between Wave and Crab environments in a single Swedish site (SWn4). Some candidate inversion regions (chiefly those in LG6, LG14 and LG17; Table S1) contained clusters of strongly clinal (putatively nonneutral) SNPs, but not all showed high levels of Crab-Wave FST differentiation (34) and most remained polymorphic in one or both habitats (24). Here, we investigated the role of inversions across a much larger geographical scale from Spain to Sweden and observed a similar pattern, with some SNPs having a strong signal of covariation with habitat contrasts (i.e. BayPass test) but low F_ST_ values and outlier sharing (Fig. 2A and D). This finding suggests that polymorphic inversions could be maintained as balanced polymorphisms that vary in equilibrium frequency according to habitat. On the other hand, the presence of chromosomal inversions can confound the perceived amount of genetic sharing we observed. For instance, we found that sharing increases when measured at different scales from SNPs to 500 bp windows to inversion regions, suggesting that larger shared genomic regions could contain many non-shared SNPs. This could be explained partly by the inherent limitation of defining outlier SNPs as those in the top 1% of the FST distribution. Moreover, it is likely that, given the recombination suppression effect imposed by chromosomal inversions, a large number of linked SNPs within the inversions are neutral hitchhikers, leading to overall low sharing at the SNP level.

Parallelism cannot be explained fully by the presence of putative chromosomal inversions, as we also found shared outlier loci scattered across the genome (cf. 9). This is consistent with polygenic selection of multiple loci of small effect underlying parallel adaptive divergence in *L. saxatilis* (33, 34, 39, 47). Our results contain an unavoidable bias because chromosomal inversions are often selected as large haplotype blocks, and thus are expected to have a strong signal of divergence in a genomic scan like ours. This confounds estimates of the contribution of loci within *versus* outside inversion regions to adaptive divergence. In addition, some clusters of shared genetic differentiation were observed in regions with no chromosomal inversions (e.g. LG2 and LG8). This may be due to the presence of other currently undetected chromosomal inversions, or due to alternative mechanisms of suppressed recombination (e.g. chromosomal centromeres; 53) in combination with background selection or the divergence of strongly selected genes and their hitchhiking neighbours (57).

### Genomic architecture regulates parallel divergence at two environmental dimensions

Our results demonstrate that considering environmental heterogeneity across multiple instances of divergence is crucial to understanding the genetic basis of parallel evolution (11, 12, 14, 15). When parallel divergence is measured using a single environmental contrast, it is possible that other axes of environmental variation can confound the estimation of genetic differentiation. Particularly, alternative axes of environmental variation that are associated with more cryptic phenotypic differences, as in the case of physiological differences between low-shore and high-shore habitat contrasts in *L. saxatilis*, are easily missed. The contribution of environmental heterogeneity has been discussed previously, but not quantified, in other widely studied systems that have focused on binary axis of parallel divergence, e.g. *Midas* cichlid fish species (58), North American lake whitefish species (59), *Timema* stick-insect ecomorphs (49) and *Pundamilia* cichlid fish species pairs (60), but see (14, 15) in stickleback fish. In *L. saxatilis*, measuring genetic differentiation without accounting for its directionality (e.g. with F_ST_) would have provided an erroneous perception of sharing due to the confounding effects of different axes of parallel divergence. Explicitly incorporating the directionality of these axes, as we did here with our DI metric, can clarify their relative contributions, providing a more accurate representation of genetic differentiation across multiple instance of parallel divergence.

In conclusion, our findings reveal that considering multiple factors is essential to understand the genetic architecture underlying parallel divergence. The demographic and geographic context, the congruence in direction between distinct environmental axes, the maintenance and re-distributing via gene flow of alleles contained within chromosomal inversion polymorphisms all play into the resulting patterns. We highlight that an approach considering all of these effects provides an important gain in inferential and explanatory power and significantly contributes to shedding light on the repeatability of evolution.

## Methods

We sequenced 22 pools of individuals, 16 including 100 female individuals and six including 24 individuals from both sexes (Table S2). Our sequencing and bioinformatics pipeline is described in Supplementary Methods. Population genomic analyses were performed with filtered files with an average coverage of 68X (min = 14X; max = 204X; sd = 18X) using Popoolation2 (61) and custom scripts. Identification of outlier loci was performed with Popoolation2 for F_ST_, BayPass (51) and custom scripts for the Cluster Separation Score and the Divergence Index metric. Customs script were written in R v3.3.2 (62). Full details for material and methods can be found in Supplementary Methods.

## Data availability

All the custom scripts can be found in the GitHub repository: https://github.com/hmoral/Ls_pool_seq. Raw sequencing reads were deposited in the Sequence Read Archive (SRA) under the BioProject PRJNA494650

## Acknowledgments

We are grateful to J. Galindo, B. Johannesson and M. Ravinet for their help collecting samples, to Z. Zagrodzka for help processing samples, to M. Gautier for his advice on BayPass and to F. Raffini and S.R. Stankowski for their comments on an earlier draft. Bioinformatics analyses were run on the Albiorix cluster (http://albiorix.bioenv.gu.se/) and we thank M. Töpel for his help. This work was supported by the Swedish Research Councils Vetenskapsràdet and Formas through a Linnaeus grant to the Centre for Marine Evolutionary Biology (217-2008-1719), the Royal Swedish Academy of Sciences (CR2015-0079), the Natural Environment Research Council (NE/K014021/1 and NE/P001610/1), the European Research Council (AdG-693030- BARRIERS) and by the Science for Life Laboratory, Swedish Genomes Program. The Swedish Genomes Program has been made available by support from the Knut and Alice Wallenberg Foundation. AMW and RF were funded by the European Union’s Horizon 2020 research and innovation programme under the Marie Sktodowska-Curie Grant Agreement No. 754411 and No. 706376, respectively.

